# Effect of number of annual rings and tree ages on genomic predictive ability for solid wood properties of Norway spruce

**DOI:** 10.1101/2020.03.26.010611

**Authors:** Linghua Zhou, Zhiqiang Chen, Lars Olsson, Thomas Grahn, Bo Karlsson, Harry X. Wu, Sven-Olof Lundqvist, María Rosario García-Gil

## Abstract

Genomic selection (GS) or genomic prediction is considered as a promising approach to accelerate tree breeding and increase genetic gain by shortening breeding cycle, but the efforts to develop routines for operational breeding are so far limited. We investigated the predictive ability (PA) of GS based on 484 progeny trees from 62 half-sib families in Norway spruce *(Picea abies* (L.) Karst.) for wood density, modulus of elasticity (MOE) and microfibril angle (MFA) measured with SilviScan, as well as for measurements on standing trees by Pilodyn and Hitman instruments. GS predictive abilities were comparable with those based on pedigree-based prediction. The highest PAs were reached with at least 80-90% of the dataset used as training set. Use of different statistical methods had no significant impact on the estimated PAs. We also compared the abilities to predict density, MFA and MOE of 19 year old trees with models trained on data from coring at different ages and to different depths into the stem. 78-95% of the maximal PAs obtained from coring to the pith at high age were reached by using data possible to obtain by drilling 3-5 rings towards the pith at tree age 10-12, thereby shortening the cycle and reducing the impact on the tree.

## Introduction

Norway spruce is one of the most important conifer species in Europe in relation to economic and ecological aspects^1^. Breeding of Norway spruce started in the 1940s with phenotypic selection of plus-trees, first in natural populations and later in even-aged plantations^2^. Norway spruce breeding cycle is approximately 25–30 years long, of which the production of seeds and the evaluation of the trees take roughly one-half of that time^3^.

Genomic prediction using genome-wide dense markers or genomic selection (GS) was first introduced by Meuwissen^4^ The method modelling the effect of large numbers of DNA markers covering the entire genome and subsequently predict the genomic value of individuals that have been genotyped, but not phenotyped. As compared to the phenotypic mass selection based on a pedigree-based relationship matrix (***A*** matrix), genomic prediction relies on constructing a marker-based relationship matrix (***G*** matrix). The superiority of the G-matrix is the result of a more precise estimation of genetic similarity based on Mendelian segregation that not only captures recent pedigree but also the historical pedigree^5–7^, and corrects possible errors in the pedigree^8,9^.

There are multiple factors affecting genomic prediction accuracy such as the extent of linkage disequilibrium (LD) between the marker loci and the quantitative trait loci (QTL), which is determined by the density of markers and the effective population size (Ne). Increased accuracy with higher marker density has been reported in simulation^10^ and empirical studies in multiple forest tree species including Norway spruce^11–14^, and SNP position showed no significant effect^15–17^ Simulation^10^ and empirical^18^ studies also agree on the need of a high marker density in populations with larger effective size (Ne) in order to cover more QTLs under low LD in contributing to the phenotypic variance.

In forest tree species the accuracy of the genomic prediction model has been mainly tested in cross-validation designs where full-sibs and/or half-sibs progenies within a single generation are subdivided into training and validation sets^10, 19–22^. Model accuracy was reported to increase with larger training to validation set ratios^11, 17, 23^, while the level of relatedness between the two sets is considered as a major factor^10, 15–17, 19, 24^ When genomic prediction is conducted across environments, the level of genotype by environment interaction (GxE) of the trait determines its efficiency^11, 20, 21, 25^. The number of families and progeny size have also been shown to affect model accuracy^11, 15^.

As compared to the previously described factors, trait heritability and specially trait genetic architecture are intrinsic characteristics to the studied trait in a given population. Those two factors can also be addressed by choosing an adequate statistical model depending on the expected distribution of the marker effects^26^. Despite theory and some results indicate that complex genetic structures obtain better fit with models that assume equal contribution of all markers to the observed variation, traits like disease-resistance are better predicted with methods where markers are assumed to have different variances^13, 20, 22, 27, 28^. However, results in forestry so far indicate that statistical models have little impact on the GS efficiency^12, 17, 29^.

In this study, we conducted a genomic prediction study for solid wood properties based on data from 23-year old trees from open-pollinated (OP) families of Norway spruce. We focused on wood density, microfibril angle (MFA) and modulus of elasticity/wood stiffness (MOE) measured both with SilviScan in the lab, on standing trees of Pilodyn penetration depth and Hitman velocity of sound. The measurement methods are detailed in the next section.

The specific aims of the study were: (i) to compare narrow-sense heritability (*h*^2^) estimation, predictive ability (PA) and prediction accuracy (PC) of the pedigree-based (ABLUP) models with marker-based models based on data from measurements with SilviScan on increment cores and from Pilodyn and Hitman measurements on standing trees, (ii) to examine the effects on model PA and PC of different training-to-validation set ratios and different statistical methods, (iii) to compare some practical alternatives to implement early training of genomic prediction model into operational breeding.

## Material and methods

### Plant material

The study was conducted on two open-pollinated (OP) progeny trials: S21F9021146 (F1146) (Höreda, Eksjö, Sweden) and S21F9021147 (F1147) (Erikstorp, Tollarp, Sweden). Both trials were established in 1990 with a spacing 1.4m×1.4m. Originally, the experiments contained more than 18 progenies from 524 families at each of site, but after thinning activities in Höreda and Erikstorp in 2010 and 2008, respectively, about 12 progenies per family were left. In 2011 and 2012, six trees per site (524 * 12 ~ 6000 trees) were phenotyped^30^. Standing tree-based measurements with Pilodyn and Hitman were performed on the same trees in 2011 and 2013, respectively, after which further thinning was performed. For this study, in 2018, we generated genomic (SNP) data from 484 remaining progeny trees after thinning which belonged to 62 of the OP families (out of the original 524 families) and on average eight progenies per family. This genotypic data was combined with available phenotypic data for the same trees that were used.

### Phenotypic data

The phenotypic data was previously described in Zhou et al, 2019^31^. Increment cores of 12mm diameter from pith to bark were collected from the progenies in 2011 and in 2012. These samples were analyzed for pith to bark variations in many woods and fiber traits with a SilviScan^32^ instrument at Innventia (now RISE), Stockholm, Sweden. This data is referred as increment core-based measurements through the text. The annual rings of all samples were identified, as well as their parts of earlywood, transition wood and latewood, averages were calculated for all rings, as well as their parts and dated with year of wood formation^33^.

The aim of breeding is not for properties of individual rings, but properties of the stem at harvesting target age. Therefore, this study focused on predictions of averages for stem crosssections, and we chose tree age 19 years as the reference age, with models trained on trait averages for all rings formed up to different younger ages. Three types of averages were calculated and predictions compared for density, MFA and MOE: 1) area-weighted averages, relating to the cross-section of the stem, 2) width-weighted, relating to a radius or an increment core, and 3) arithmetic averages, where all ring averages are weighted with same weight. For the calculation of area-weighted average we assumed that each growth ring is a circular around the pith, calculated the area of each annual ring from its inner and outer radii, and when calculating the average at a certain age, the trait average for each ring was weighted with the ring’s proportion of the total cross-sectional area at that age. Similarly, for the calculation of the width-weighted average, the trait average for each ring was weighted with the ring’s proportion of the total radius from pith to bark at that age. Similar results were obtained with the three average methods. For this reason, only the estimates based on the area-weighted method (the most relevant for breeding) are shown. Tree age 19 years was used as the reference age. Thus, all the selection methods investigated for density, MFA and MOE, phenotypic and genetic, were compared based on how well they predicted the cross-sectional averages of the trees at this age, with their last ring formed during the vegetation period of 2009.

In addition, estimates of the three solid wood traits were calculated based on data from Pilodyn and Hitman instruments, measured on the standing trees without removing the bark at age 22 and age 24 years, respectively. Pilodyn measures the penetration depth with a needle pressed into the stem, which is inversely correlated with wood density. Hitman measures the velocity of sound in the stem, which correlates with microfibril angle, MFA^34, 35^. MOE is related to wood density and velocity of sound^36–38^ and can therefore be estimated by combining the Pilodyn and Velocity data, which estimates we here name MOE_ind_ (for standing-tree based). Further details on how this was performed in our study are given in Chen et. al 2015^39^. The references show that these standing-tree-based measurements provide useful information and are very time and cost-efficient. However, they do not allow calculation of properties of the tree at younger ages. Therefore, we were not able to investigate from what early ages such data can be uses within genomic selection.

### Genotypic data

Genomic DNA was extracted from buds or needles when buds were not available. Qiagen Plant DNA extraction protocol was utilized for DNA extraction and purification and DNA quantification performed using the Qubit^®^ ds DNA Broad Range (BR) Assay Kit (Oregon, USA). Genotyping was conducted at Rapid Genomics, USA, using exom capture methodology same as the method used in Baison et. al 2019^40^. Sequence capture was performed using the 40 018 diploid probes previously designed and evaluated for *P. abies*^41^ and samples were sequenced to an average depth of 15x using an Illumina HiSeq 2500 (San Diego, USA)^40^. Variant calling was performed using the Genome Analysis Toolkit (GATK) HaplotypeCaller v3.6^42^ in Genome Variant Call Format (gVCF) output format. After that, the following steps were performed for filtering: 1) removing indels; 2) keeping only biallelic loci; 3) removing variant call rate (“missingness”) < 90%; 4) removing minor allele frequency (MAF)<0.01. Beagle v4.0^43^ was used for missing data imputation. After these steps, 130,269 SNPs were used for downstream analysis.

### Population structure

As a first step, we conducted a principal component analysis to determine the presence of structure in our population. The spectral decomposition of the marker matrix revealed that only about 2% of the variation was captured by the first eigenvector, indicating low population structure. Additionally, in previous study, low genotype by environment (GxE) interaction was detected for wood quality traits on these two trials^30^. Therefore, population structure was not considered in the design of cross-validation sets (see Modelling and cross-validation chapter for further details on the cross-validation sets design).

### Narrow-sense heritability (h^2^) estimation

For each trait, an individual tree model was fitted in order to estimate additive variance and breeding values:

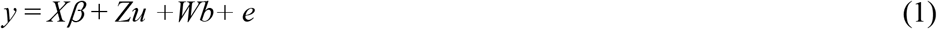

where *y* is a vector of measured data of a single trait, *β* is a vector of fixed effects including a grand mean, provenance and site effect, *b* is a vector of post-block effects and *u* is a vector of random additive (family) effects which follow a normal distribution *u*~N(0,Aσ^2^_u_) and *e* is the error term with normal distribution N(0,Iσ^2^_e_). *X, Z* and *W* are incidence matrices, A is the additive genetic relationship matrix and I is the identity matrix. σ^2^_u_ equals to σ_a_^2^ (pedigree-based additive variance) when random effect in equation 1 is pedigree-based in which case *u*~N(0,Aσ^2^_u_), and σ^2^_u_ equals to σ_g_^2^ (marker-based additive variance) when random effect in equation 1 is marker-based in which case *u*~N(0,Gσ^2^_u_). The G matrix is calculated as *G* = 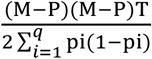, where M is the matrix of samples with SNPs encoded as 0, 1, 2 (i.e., the number of minor alleles), P is the matrix of allele frequencies with the ith column given by 2(pi – 0.5), where pi is the observed allele frequency of all genotyped samples.

Pedigree-based individual narrow-sense heritability (*h_a_*^2^) and marker-based individual narrow-sense heritability (h_g_^2^) were calculated as

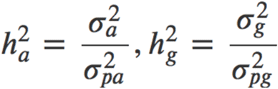

respectively, *σ*^2^_*pa*_ and *σ*^2^_*pg*_ are phenotypic variances for pedigree-based and marker-based models, respectively.

### Selection of the optimal training and validation sets ratio

Cross-validation was conducted after dividing randomly the whole dataset into a training and a validation set. To find the most suitable ratio between the two, we divided the data into sets with five different ratios between the training and the validation sets: 50, 60, 70, 80 and 90%. 100 replicate iterations were carried out for each tested ratio and trait.

### Statistical method for model development

In the same context we aimed to find optimal methods. Several statistical methods were compared: pedigree-based best linear unbiased predictions (ABLUP), and four GS methods: genomic best linear unbiased predictions (GBLUP)^44^, random regression-best linear unbiased predictions (rrBLUP)^4, 45^, BayesB^46^, and reproducing kernel Hilbert space (RKHS). rrBLUP used a shrinkage parameter lamda in a mixed model and assumes that all markers have a common variance. In BayesB the assumption of common variance across marker effects was relaxed by adding more flexibility in the model. RKHS does not assume linearity so it could potentially capture nonadditive relationships^47^ R package *rrBLUP*^48^ was used for GBLUP and rrBLUP, package *BGLR*^49^ was used for BayesB and RKHS. The pedigree-based relationship matrix was obtained with the R package *pedigree* (Coster2013).

### PA and accuracy estimation

The adjusted phenotypes y’=y-*Xβ* were used as model response in the genomic prediction models. Model quality was evaluated by predictive ability (PA), which is the mean of the correlation between the adjusted phenotype and the model predicted phenotypes, r(y’,yhat) from 100 times CV. Prediction accuracy (PC) was defined as PA/ √(h^2^)^15, 50^. In order to investigate whether GS model training can be conducted at earlier age, PA at each tree calendar age and cambial age were estimated. In this case, cross validation was conducted only using area-weighted values at each age, then the trait values at each age were estimated. PA at a specific age was calculated as the correlation between estimated trait values at that age and area-weighted values from pith to the last ring (for cambial age) and last year (for calendar age), respectively.

Genomic selection for well-performing trees with the use of marker information (G matrix) requires access to previously trained GS models. Thus, model training is a necessary part of GS integration into operational breeding. Model training can be conducted in already existing plantations with trees of relatively high ages, as illustrated in this work. It is, however, expected and desired that such model training can be conducted with high PAs also for younger trees. This would be especially useful if maturity (flower production) can be accelerated, to shorten the total breeding cycle.

Operationally, it is also important to develop protocols to assess wood quality in resources at minimum cost and time, and with minimal impact on the trees. Therefore, on coring, it is not only important to know the minimum age at which useful information can be obtained, but also from how many rings from the bark towards the pith information is required to train models with high predictive ability. To address these two practical questions for operational breeding, we trained prediction models based on data from different sets of rings, in order to mimic and compare PAs obtained when coring at different ages of the trees to different depths into the stem, or more precisely, using data from different numbers of rings, starting next to the bark. All the models were judged on, compared by their ability to predict the cross-sectional average of the trait at age 19 years across all trees in the validation set.

## Result

### Narrow-sense heritability (h^2^) of the phenotypic traits, predictive ability (PA) and Predictive accuracy (PC) based on pedigree and maker data

In Table 1, narrow sense heritabilities (*h*^2^) and Prediction Abilities (PA) based on ABLUP and GBLUP are compared for density, MFA and MOE based on cross-sectional averages at age 19 years, and for Pilodyn, Velocity and MOE_ind_ based on measurements with the bark at age 22 and 24 years, respectively. For density, MOE and Pilodyn, *h*^2^ did not differ significantly between estimates based on the pedigree (ABLUP) and marker-based (GBLUP) methods taking standard error into account. For MFA, the pedigree-based *h*^2^ was lower than the GBLUP estimate while for Velocity and MOE_ind_, the pedigree-based *h*^2^ was higher.

**Table 1.** Trait heritability, predictive ability (PA) and predictive accuracy (PC) Predictive accuracy (PC)for density, MFA and MOE cross-sectional averages at tree age 19 years, for their proxies on the stems without removing the bark at tree ages 21 and 22 years. Standard errors are shown in within parenthesis.

| Trait | Narrow-sense heritability<br>(standard error)<br>($h^2$) | | Predictive ability<br>(standard error)<br>(PA) | | Predictive Accuracy<br>(PA/h) | |
| --- | --- | --- | --- | --- | --- | --- |
|  | ABLUP | GBLUP | ABLUP | GBLUP | ABLUP | GBLUP |
| density | 0.70 (0.18) | 0.69 (0.15) | 0.30 (0.01) | 0.29 (0.03) | 0.36 | 0.35 |
| MFA | 0.04 (0.08) | 0.17 (0.13) | 0.04 (0.01) | 0.16 (0.02) | 0.20 | 0.39 |
| MOE | 0.27 (0.14) | 0.31 (0.15) | 0.15 (0.01) | 0.22 (0.02) | 0.29 | 0.39 |
| Pilodyn | 0.35 (0.15) | 0.32 (0.14) | 0.22 (0.01) | 0.20 (0.01) | 0.37 | 0.35 |
| Velocity | 0.16 (0.12) | 0.11 (0.10) | 0.10 (0.01) | 0.13 (0.01) | 0.25 | 0.39 |
| MOEind | 0.31(0.14) | 0.17 (0.13) | 0.17 (0.01) | 0.19 (0.01) | 0.31 | 0.46 |

ABLUP=pedigree-based Best Linear Unbiased Predictor (BLUP); GBLUP= genomic-based BLUP.

When using pedigree, the order of the traits by *h*^2^ agrees with their order by PA estimates. Traits with higher *h*^2^ tended to show also high PA estimates irrespective of the method. The ABLUP PA estimates were similar to the GBLUP estimates for density and Pilodyn, while for the rest of the traits GBLUP delivered slightly higher PA estimates, and significantly higher for MFA. The relative performances of ABLUP compared to GBLUP differed for MOE, Velocity and MOE_ind_. The *h*^2^ estimates for MOE were similar for both methods, while the PA estimate was higher for GBLUP. In the case of Velocity and MOE_ind_, a higher *h*^2^ based on pedigree contrasted with a slightly higher PA estimates based on marker data. Standardization of the PAs with the h values did not change the conclusions on the relative efficiencies of pedigree versus marker data-based estimates.

### Marker-based PA and PC between increment core-based and standing-base wood quality traits

The marker-based PAs were generally 25-30% higher for traits density, MFA and MOE measured with SilviScan than for their respective standing tree-based method which measured with Pilodyn and Hitman. Concordantly, the *h*^2^ values were 46%, 65% and 55% higher based on Silviscan methods, respectively. However, if we compare PC of the increment core- and standing tree-based methods, they were similar, and PC of MOE_ind_ was even higher than that for MOE using GBLUP.

### Effects on PAs of the GS models ratios between the training and validation sets, and from the statistical method used

Figure 1 shows how the PA estimates change with increasing percentage of data used for training of the GS model (training set), and as a consequence decreasing validation set, on use of the five studied statistical methods: one based on pedigree data and four on marker information. For most of the traits, PA estimates showed a moderate increase with increasing training set, irrespective of the statistical method. Exceptions were observed for MFA and MOE with less clear trends and the highest PA estimates at 80% of the trees in the training set. Figure 1 also shows that the PAs were consistently about 25-30% higher for density, MFA and MOE compared to their proxies-based om measurements with Pilodyn and Hitman: approximately 0. 28 versus 0.18, 0.17 versus 0.13 and 0.25 versus 0.18, respectively.

**Figure 1.**
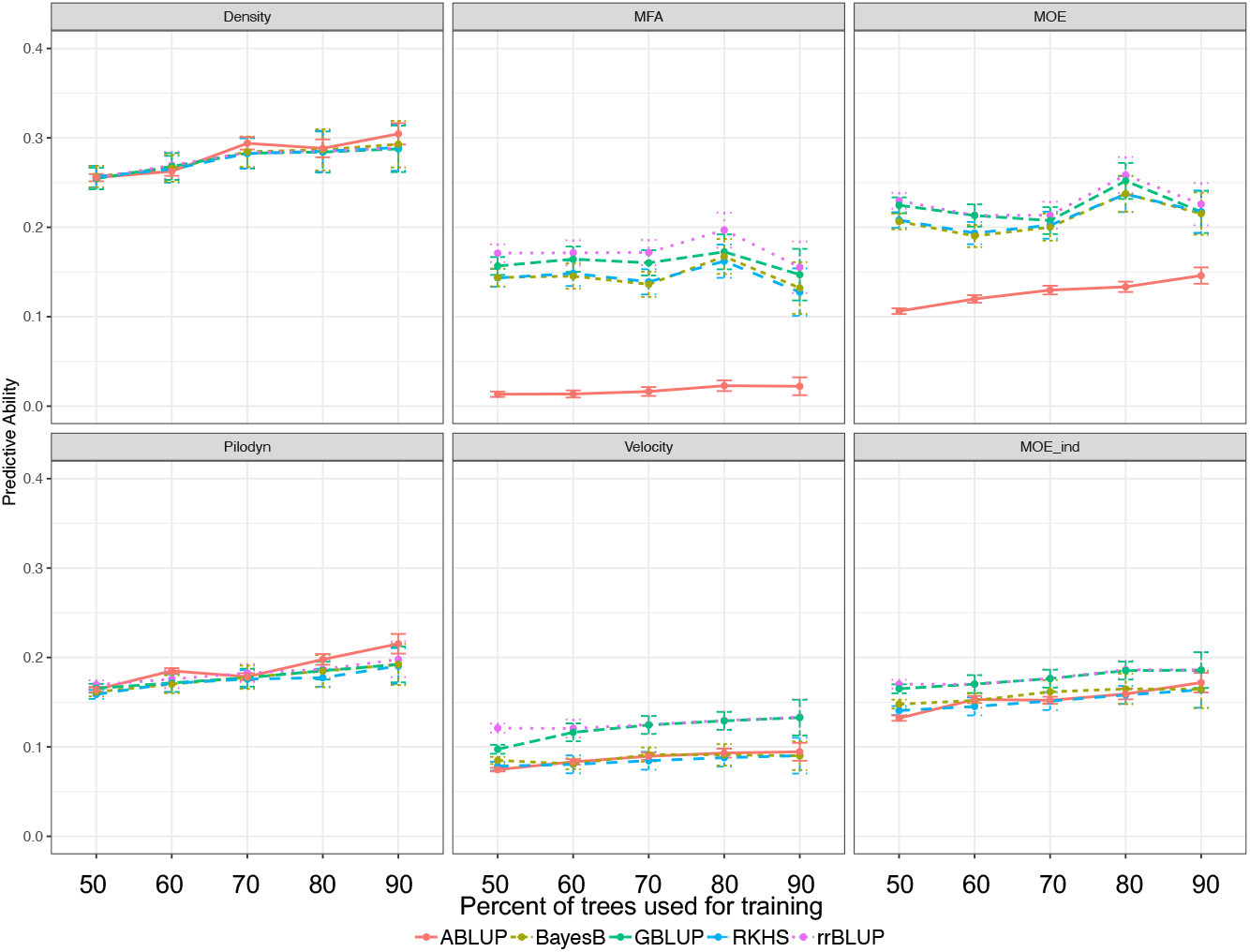
Predictive ability obtained with different ratios of training set and validation set, using different statistical methods.

For density and Pilodyn, all five methods resulted in very similar PA estimates across the ratios, while rrBLUP and GBLUP seemed superior for the rest of the traits, and mostly so for Velocity and MOE (Figure 1). The rest of the analysis were conducted based on the GBLUP modelling method.

### PAs on estimation of traits at reference age with models trained on data available at earlier ages

Figure 2 shows how well the cross-sectional averages of the different traits at the reference age 19 years were predicted by models trained based on data from the rings between pith and bark at increasing ages, using the GBLUP method. The calculations were performed with two representations of age: 1) Tree age counted from the establishment of the trial (calendar age) and 2) cambial age (ring number). In a plantation, the tree age of a planted tree is normally known but not the cambial age at breast height, as it depends on when the tree reached the breast height. For the trees originally accessed, almost 6000 trees from the two trials, this age ranged from tree age 2 to 15 years^33^. Among the 484 trees investigated in the current study, only 60 trees representing 33 families had reached breast height at tree age 3 years, 248 trees at 4 years and 410 at age 5 years (Figure 2). This means that for tree age, data are only available from year 3, and then for only 12% of the trees. Those trees being identified based on fast longitudinal growth but also typically fast-growing radially. It was previously described a positive correlation of R^2^ = 0.67 familywise between radial and height grown across almost 6000 trees (Lundqvist at al 2018). Thereafter, the number of trees increased and reached the full number some years later. When studying the trees based on cambial age, the pattern is adverse with data for all trees at ring 1 but decreasing numbers when approaching the tree age of sampling. The number of trees included in this work at each tree and cambial age are shown with grey bars in Figure 2.

**Figure 2.**
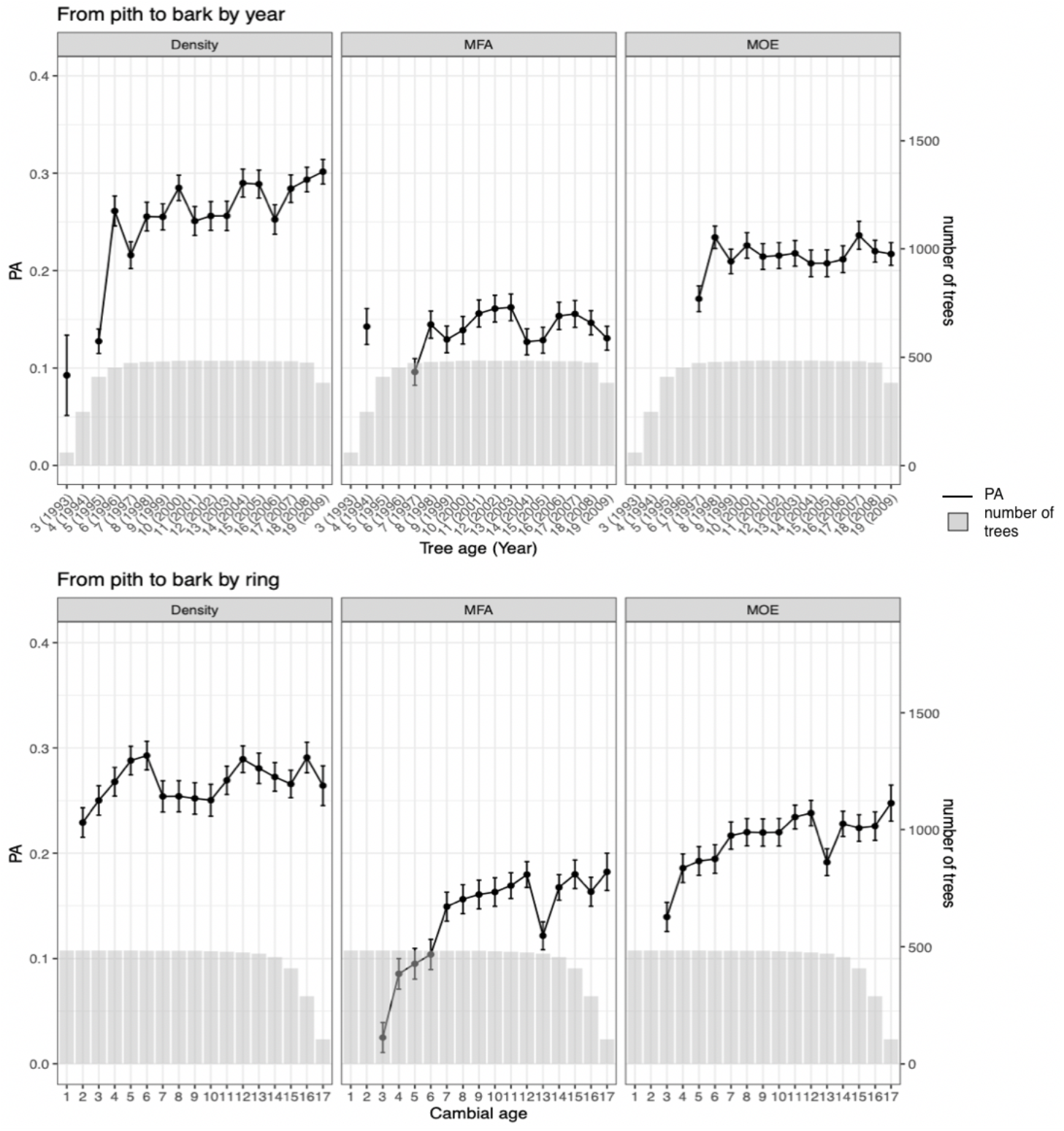
Estimated Predictive abilities (PA) for prediction of cross-sectional averages at tree age 19 years, based on cross-sectional averages at different tree ages (upper graphs) and cambial ages (lower graphs) from pith to bark.

For density, the estimated PAs showed a rising trend within a span of about 0.25-0.30 for the models based on both age types, after the first years. But the year-to-year fluctuations were more intense for models based on data organized on tree age. As MFA typically develops from high values at the lowest cambial ages via a rapid decrease to lower and more stable values from cambial age 8-12 years and on, one may expect that models trained on data from only low ages would have difficulties to predict properties at age 19 years. This was also confirmed. We even obtained some negative PA values at early ages, such as years 1995 and 1996, and the PAs for cambial age-based models started from very low values, then increasing. The curves for MOE showed PAs developing at values in between those for density and MFA. This is logical, as MOE is influenced by both density and MFA, with particularly negative effects from the high MFAs at low cambial ages. At cambial age 13, MFA and MOE showed a drop in the cambial age-based PA estimates. Generally, the Figure indicates that genomic selection for density could be conducted at an earlier age than for MFA and MOE.

### Search for optimal sampling and data for training of GS prediction models

Figure 2 showed estimated PAs of models trained on data from sampling different years, using data from all rings available at that age (except for the innermost ring). In this section instead of estimating PAs with the whole increment core from bark to pith, we estimated PAs with partial cores with different shorter depths to reduce the injury to the tree, as showed in Figure 3a-d. This analysis was preformed based on tree age data only, as the cambial age of a ring can only be precisely known if the core is drilled to the pith which allowing all rings to be counted.

**Figure 3.**
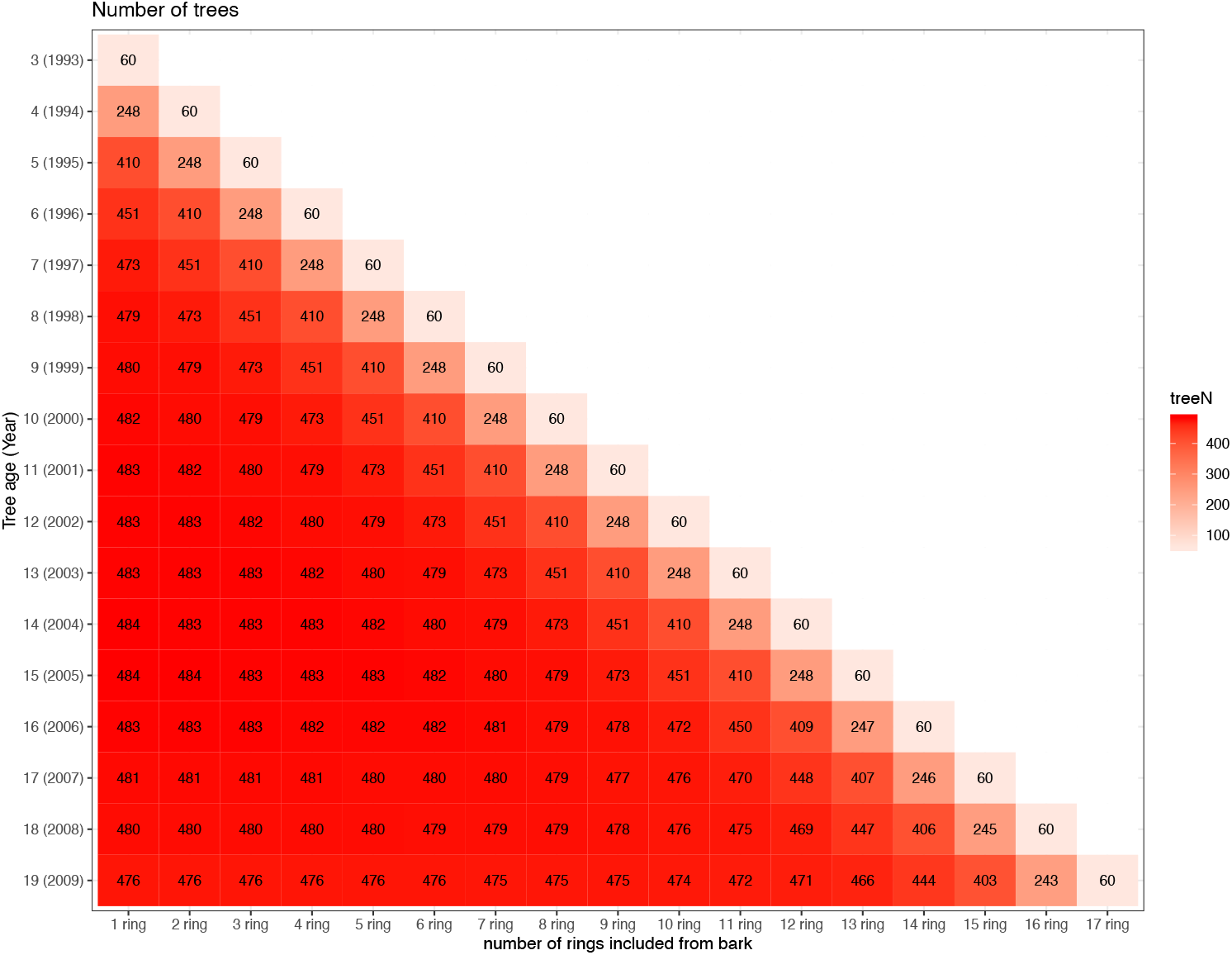
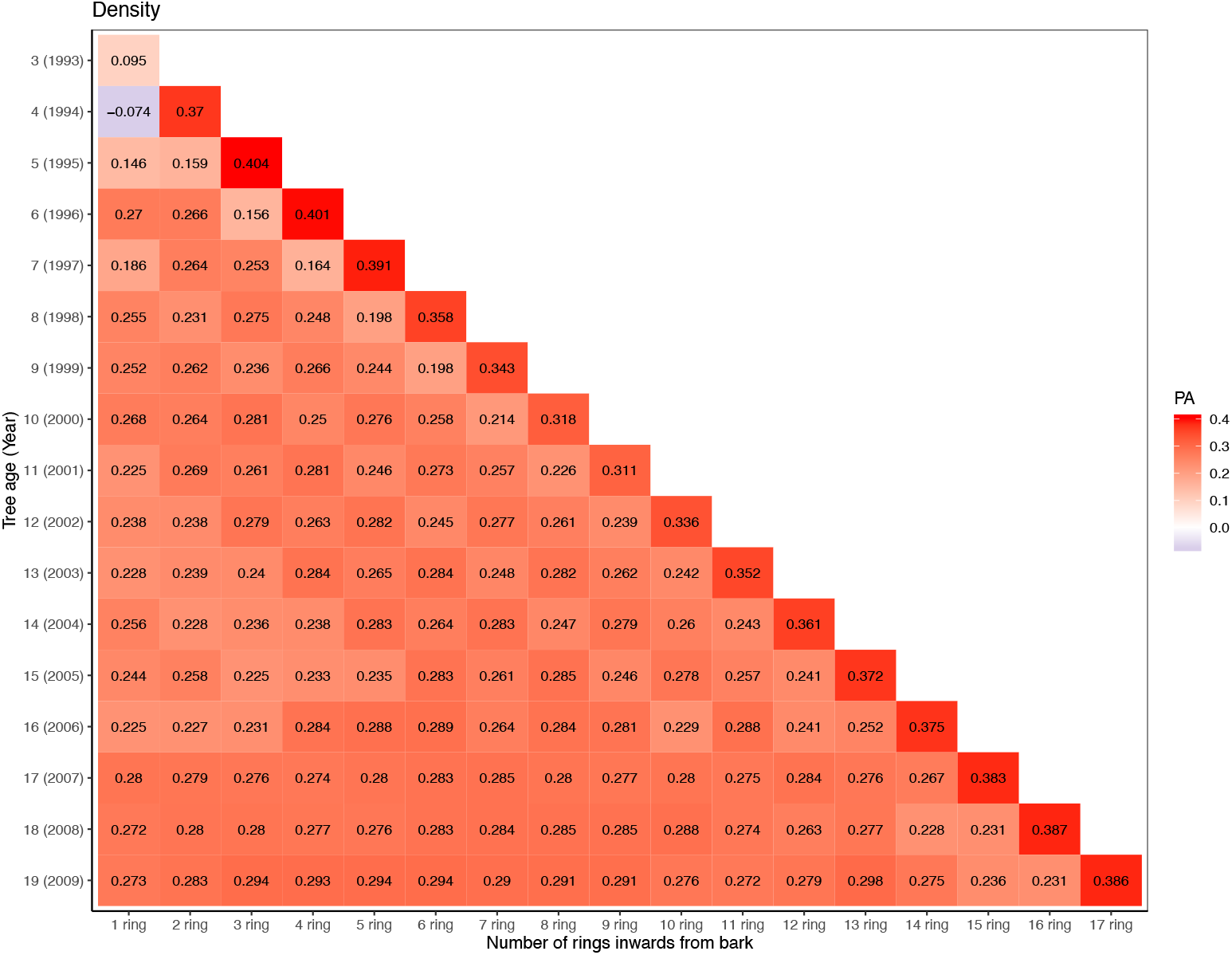
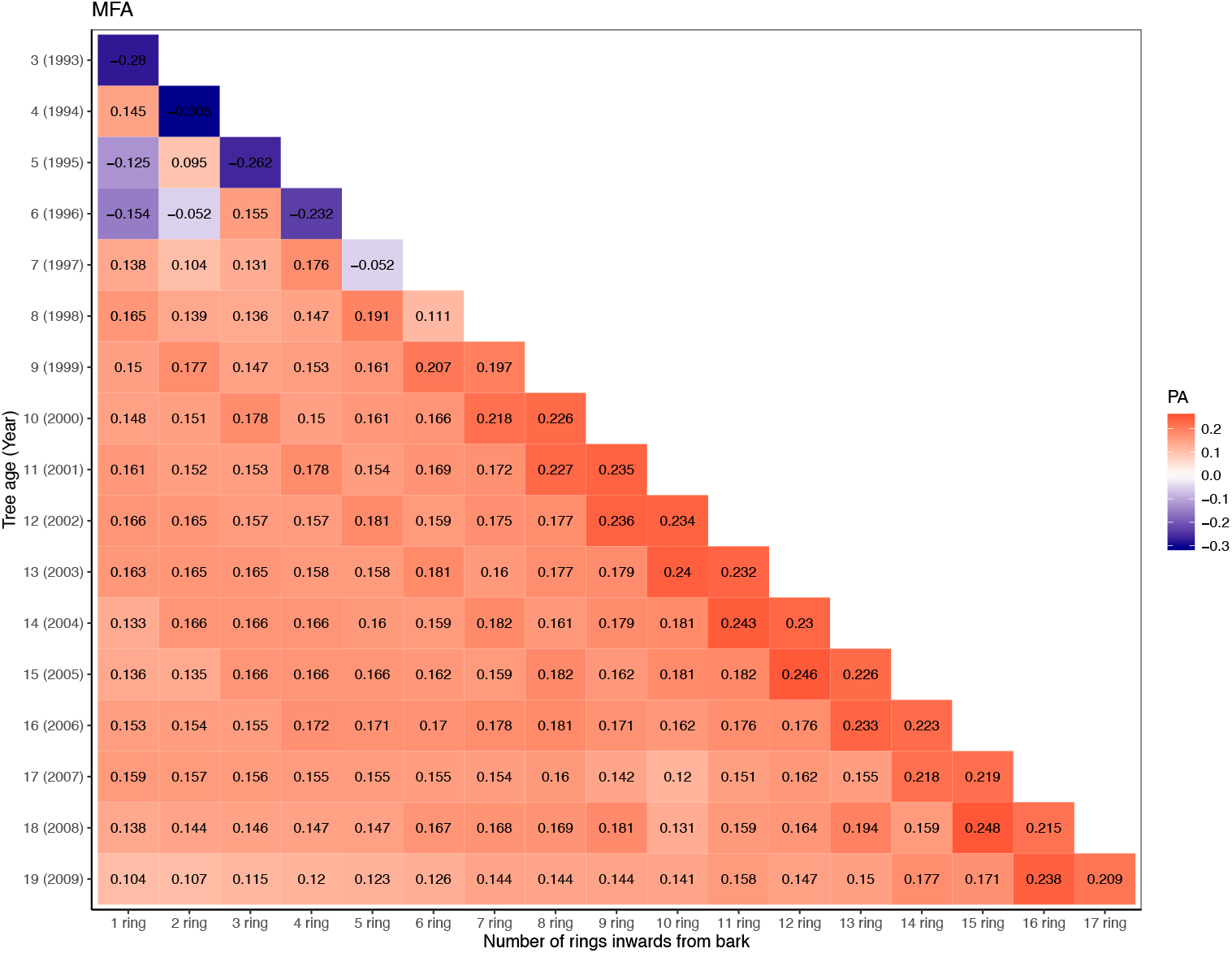
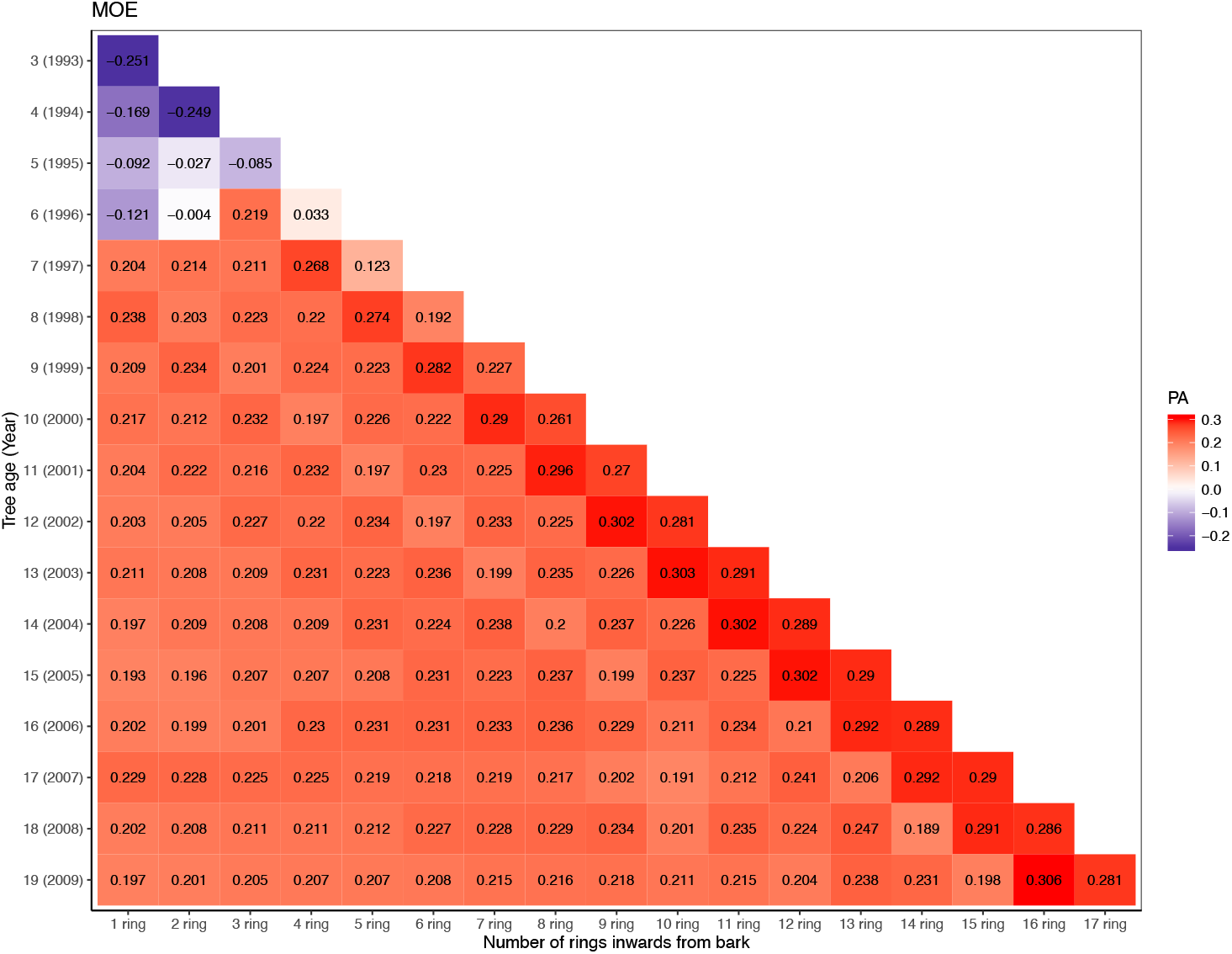
Predictive ability from bark to pith at different tree ages (y-axis) and an increasing number of rings included in the estimation (x-axis). 3a) Number of trees at each tree age with different number of rings. 3b) PA of density at each tree age with different number of rings. 3c) PA of MFA at each tree age with different number of rings 3d) PA of MOE at each tree age with different number of rings.

Each row of the figures represents a tree age when cores are samples, starting at age 3 years when the first 60 trees formed a ring at breast height, ending at the bottom with the reference age 19 years with17 rings. Each column represents a depth of coring, counted in numbers of rings. As one more ring is added each year, thus also to the maximum possible depth on coring, the tables are diagonal. The uppermost diagonal represents models trained on data from the 60 (12%) trees which had reached breast height at age 3. The diagonal next below represents models based on the 243 (51%) trees with rings at age 4, etc. The PAs shown below the three uppermost diagonals represent models trained of data from more than 90% of the trees. The PAs were calculated from the cross-validation, based on data from the trees on which the respective models were trained. This means that the PAs of the three uppermost diagonals are based only on fast-growing trees not fully representative for the trials. Many of the highest PAs found occur along these diagonals. Due to their trees’ special growth, only PAs based on more than 90% of the trees will be further commented.

For wood density, Figure 3b, the variations in predictability show an expected general pattern: The PAs increased with the increase of tree age on coring, and also with the increase of depth, the increase of number of rings from which the cross-sectional averages were calculated and exploited on training of the prediction models. The highest values, 0.29, are obtained at age 19 years, but then also data from the reference year are included on training the prediction model. An example of quite high PAs at lower ages and depths: For coring at tree ages 10-12 years and using data from the 3-5 outermost rings, all alternatives gave PA values of 0.26-0.29.

For MFA, a trait with low heritability, the PA values are low as already shown in Figure 2 and the pattern in Figure 3c is not easy to interpret. Here, the same set of alternatives of samples at tree ages 10-12 and depths 3-5 outermost rings gave PA values of 0.15-0.18, compared to the maximum of 0.19 among all alternatives using 90% of the trees. The values are lower at the highest ages. Streaks of higher and lower values can be imagined along the diagonals. The pattern for MOE in Figure 3d is similar to that of MFA, but on higher level. Training on data from coring at ages and to depths as above gave PA values of 0.20-0.23, compared to the corresponding maximum of 0.25.

## Discussion

We have conducted a genomic prediction study for solid wood properties assessed on increment cores from Norway spruce trees with SilviScan derived data from pith to bark, using properties of annual rings formed up to tree age 19 years as the reference age.

On Norway spruce operational breeding, the use of OP families is preferable because it does not require expensive control crosses. The only action required is to collect cones where progenies are typically assumed to be half-sibs. Thus, OP families permit evaluation of large numbers of trees at lower costs and efforts than structured crossing designs. We investigated narrow-sense heritability estimation with ABLUP and marker-based GBLUP and the effect on PA from using different training-to-validation set ratios, as well as different statistical methods. Further, we investigated what level of precision can be reached when training the models with data from trees of different ages, also compared results for the solid wood properties with those for their proxies. We also estimated the level of PAs reached when coring to different depths from the bark at different tree ages, in order to find cost-effective methods for GS with minimum impact on the trees on the acquisition of data for training the prediction models.

### Narrow-sense heritability (h^2^)

In our study, PA estimates for both pedigree and marker-based methods were consistent with their respective h^2^ estimates. A conifer literature review indicates that the level of consistency varies across studies^8, 18–20^. In our study, h^2^ estimation of density, MOE and Pilodyn were similar for ABLUP and GBLUP; for Velocity and MOE_ind_, ABLUP had higher h^2^ estimation and for MFA, GBLUP achieved higher h^2^ estimation. In a previous study conducted on full-sib progenies in Norway spruce, however, the ABLUP-based h^2^ were reported higher in all three standing-tree-based measurements^11^. Instead, other conifer studies based on full- or half-sib progenies reported a comparable performance of A-matrix and G-matrix based methods in Pinus taeda^18, 23^, Douglas-fir^29^ and Picea mariana^15^ for growth related traits and wood properties. Moreover, ABLUP accuracies were lower for growth, form and wood quality in Eucalyptus nitens^24^ Experimental design factors such as number of progenies and their level of coancestry, statistical method and the traits and pedigree errors under study may account for the apparent inconsistence in the relative performance of both methods^51^.

Our results indicate that for more heritable traits ABLUP and GBLUP capture similar levels of additive variance, whereas for traits with very low heritability using ABLUP, such as MFA, the markers are able to capture additional genetic variance probably in the form of historical pedigree reflected in the G matrix. Less obvious is the case for Velocity and MOE_ind_ where GBLUP seems to capture lower values of additive variance. It is possible that at intermediate values of *h*^2^ the benefits of capturing historical consanguinity is overcome by possible confounding effects caused by markers which are identical by state (IBS) or simply due to genotyping errors. The *h*^2^ values obtained with ABLUP and GBLUP is the result of a balance between multiple factors such as the genetic structure of the trait, the historical pedigree, and the possible model overfitting to spurious effects or genotyping errors.

### Effects on GS model predictive ability (PA) of training-to-validation sets ratios and statistical methods

In conifers and *Eucalyptus* cross-validation is often performed on 9/1 training to validation sets ratio^8, 12, 15, 16, 28^. This coincide with the general conclusion from the present study, with exception for MFA and MOE, for which the best results were obtained at ratio 8/2. It has been suggested that when the trait has large standard deviation, more training data is needed to cover the variance in order to get high predictive ability^52^. So, for density, Pilodyn and Velocity, PA kept increasing with the size of the training set increased. But for other traits with smaller standard deviation, (4.44 and 2.28 for MFA and MOE), PA decreased when increasing the training set from 80% to 90%, which may indicate that too much noise was introduced during model training.

The fact that the estimated PAs for all the solid wood properties as measured by SiliviScan are 25-30% higher than their proxies estimated from measurements of penetration depths and sound velocity at the bark may reflect the indirect nature of their proxies: the correlations calculated for the almost 6000 trees initially sampled were −0.62 between Pilodyn and density, −0.4 between Velocity and MFA and 0.53 between MOE_ind_ and MOE^39^.

In the conifer literature it has more often been reported similar performance of different marker-based statistical models for wood properties^11, 12, 18, 28, 53^. This general conclusion agrees with our findings for all our traits with the exception of Velocity and to a less extent of MOE_ind_. For these two traits, GBLUP and rrBLUP performed better than the other GS methods, which could be the result of a highly complex genetic structure where a large number of genes of similar and low effect are responsible for controlling of the trait. For traits affected by major genes the variable selection methods, for example BayesB or LASSO, have been reported to perform better^18^, whereas for additive traits the use of nonparametric models may not yield the expected accuracy^54^.

### Comparison of PA and PC from methods based on pedigree and markers

Generally, pedigree-based PA estimates in conifer species have been reported to be higher or comparable to marker-based models^11, 15, 16, 19, 20, 23^, but there are also some studies reporting marker-based PA estimates to be higher^13, 24, 55^. Our results for density and Pilodyn follow the general finding in forest trees, whereas for MFA, a low heritability trait, the PA estimation based on GBLUP model is substantially higher (0.16) compared to the ABLUP model (0.04). When PA is standardized with h, the predictive accuracies of the methods become more similar across traits, indicating that proportionally similar response to GS can be expected for all traits.

### Use of tree age versus cambial age (ring number)

From a quick look at Figure 2, one may get the impression that breeding based on cambial age data allows earlier selection than using tree age data. That would however be a too rushed conclusion. At tree age 3 years, after the vegetation period of 1993, only 12.5% of the trees had formed the first annual ring at breast height. Not until tree age 6 years, more than 90% of the trees had done so. But if aiming for 90% representation, one must wait several years more till more rings were formed at breast height, i.e., from 1993 to end of growth season 1996 at tree age 6. And to train models based on data from 90% of the trees for cambial age say, age 6 at breast height, samples cannot be collected until the end of growth season at tree age 11 years, or if a representation of 80% is judged as satisfactory, at tree age 10 years. This has to be considered if selection efficiencies are calculated based on cambial age data, which is common. Such results have for instance been published based on the almost 6000 trees sampled at 2011 and 2012^30^.

Correctly compared based on minimum 90% of the trees, the estimated PAs shown in Figure 2 are similar between the age alternatives, or slightly better for use of tree age. For example, the PA for MOE using cambial age data shows a smooth increase, reaching above 0.2 at cambial age (ring number) 7, which needs data from the tree of age 12. The corresponding curve from using tree age passed above 0.2 already at age 8 years. However, curves based on tree age often show larger year-to-year variation. This is most likely an effect of the fact that the rings of same cambial age represent wood formed across a span of years with different weather. Thus, cambial age data reflect annual weather across a range of years, which does not happen when using tree age data. On the other hand, from a practical point of view, methods based on using tree age may be easier to apply in operational breeding, especially as light color results in Figure 3b–3d, indicating that high PAs can be reached without coring all the way to the pith. To number the rings for precise cambial age, you need to find the innermost ring at the pith, but that may not be necessary for good results.

### Implementation of GS for solid wood into operational breeding

The results indicate that GS can result in similar early selection efficiency or even higher than traditional pedigree-based breeding and offers further possibilities. Previously, in loblolly pine it was reported that models developed for diameter at breast height (DBH) and height with data collected on 1 to 4-year old trees had limited accuracy in predicting phenotypes at age 6-year old^21^. In British Columbia Interior spruce, the predictive accuracy for tree height of models trained at ages 3 to 40 years, at certain intervals, and validated at 40 years revealed less opportunities for early model training, since the plateau was not reached until 30 years^28^.

In our study, the highest PA values (on the diagonals in Figure 3b–3d) were obtained for the subsets of fast-growing trees which had reached breast height already at tree age 3 and 4 years, 12% and 51% of the total number of trees, representing a limited number of the OP families included in the analysis. Trees in this subgroup are affected by high intensity of selection for alleles accelerating growth within each OP family. Also, on cross-validation the prediction abilities for this group were calculated based on the trees within the same group. In this elite group different factors could account for a higher PA value, such as lower phenotypic variance, decreased number of alleles of minor effect could also facilitate identification of major effects and/or higher consanguinity between those families which may share alleles for growth. These models are shown for completeness, but as they cannot be used for operational breeding they are not further discussed.

Models for genetic selection are useful in different steps of a breeding program. One type of prediction models, here illustrated with Table 1, can be trained from existing trials, preferably based on trees of as old age as available. Since the aim of breeding is to predict tree qualities at age of harvesting when the major part of the stem will be dominated by mature wood. Training the models in older trees for wood properties also allows considering other properties which cannot be easily observed from trees of very young age, such as stem straightness and health. For wood density, the results indicate that models can be built without coring very deep into the stem. It may be expected that this is valid also for instance for tracheid dimensions which in combination determines the wood density^33^.

As illustrated in this work, two aspects of incorporating wood properties into operational GS breeding programs can be addressed with the same set of data. Firstly, as mentioned above, models for cost-effective selection based on genomic information from existing trees. In that case, models from data at old ages would normally be preferred, for example for wood density some model at bottom line of Figure 3b. Secondly, models providing guidance on at what age it is reasonable to approach young trees for training of GS models for specific traits: a) trees in existing juvenile trials, or b) trees of new generations with different pools of genetics. As an example, the same Figure 3b for wood density suggests GS model training at tree ages 10 to 12 on the third to fifth outermost rings to reduce costs and the negative impact on the tree.

## Conclusions

1. In comparison with phenotypic selection, Genomic selection methods showed similar to higher prediction abilities (PAs) for both increment core- and standing tree-based phenotyping methods. This indicates that the standing tree-based measurements may be a cost-effective alternative method for GS, but higher PAs were obtained based on increment core-based wood analyses.
2. Different genomic prediction statistical methods provided similar PA. At least 80% data should be included in the training set in order to reach the highest levels of PA
3. This study represents the first published investigation of the efficiency of GS with prediction models trained on data acquired from sampling/coring trees at different ages, combined with sampling/coring to different depths, to optimize the operational breeding for the combination of length of breeding cycle, cost and impact on the trees. The results indicate that similar efficiency can be obtained at tree age 10-12 with 3-5 outermost rings.

## Contributions

LZ analysed data and drafted the manuscript. ZC designed sampling strategy, coordinated field sampling and edited the manuscript. BK participated in the selection of the breeding populations, providing access to field experiments and edited the manuscript. LO, TG conducted the SilviScan measurements and performed the evaluations prior to the genetic analyses. HW conceived and designed the study and edited manuscript. SOL and RRG provided ideas and revised manuscript. All authors read and approved the final manuscript.

## Acknowledgements

We would like to acknowledge the UPSC Vinnova Center of Forest Biotechnology. We also acknowledge the Swedish Research Program Bio4Energy, the Swedish Foundation for Strategic Research (SSF) and RISE for their support in phenotypic and genotypic data collection.

## Ethics approval and consent to participate

The plant materials analysed for this study comes from common garden experiments that were established and maintained by the Forestry Research Institute of Sweden (Skogforsk) for breeding selections and research purposes. Three tree breeders in Sweden were co-authors in this paper. They agreed to access the materials.

## Consent for publication

Not applicable.

